# High-bandwidth, low-profile, long-term wireless EEG telemetry allows for optogenetic entrainment of natural cortical oscillations in freely-moving rats

**DOI:** 10.1101/2024.06.05.597544

**Authors:** Beth Rees, Phillip R. Griffiths, Gavin Woodhall, Stuart D. Greenhill

**Affiliations:** Aston Institute of Health and Neurodevelopment, Aston University, Birmingham UK; AD Instruments Ltd, Oxford UK

## Abstract

Recording of whole-brain or multi-unit neuronal activity in the rodent brain is a powerful and widely used technique in neuroscience research. However, the acquisition of data from freely-moving animals is subject to a range of compromises. If a high bandwidth of data digitisation is needed, animals will either need to be tethered to the acquisition system or any telemetry used will have a short working battery life. For freely-moving experiments, especially those requiring careful behavioural measurements, such tethers and/or headstages incorporating e.g. optogenetic stimulation systems may prove to be confounding or limiting in the experiments which may be performed. Here we present the refinement and deployment of a wirelessly-charged, self-contained EEG telemeter at high data bandwidths (2kHz) with integrated optogenetic stimulator (473nm) and fully subcutaneous fibre routing and implantation. This approach has allowed for rats to be recorded long-term (6 months) without requiring device explants, charging or maintenance, with an outward appearance identical to an unimplanted rodent. We have demonstrated the use of this system to stimulate cortical networks at a range of frequencies in freely-moving and acutely-anaesthetised rats allowing for the boosting or entrainment of physiological oscillations at will.

## Introduction

The use of the electroencephalogram (EEG) and other such recordings of neuronal ensemble activity has been a prevalent and vital mechanism for investigating the function and dysfunction of the mammalian brain for much of the last century, since the inception of the EEG technique by Hans Berger in 1924. The capturing of both action potentials and subthreshold activity *in vitro* and *in vivo* have been invaluable(*1*) in our understanding of how individual cells and networks at multiple levels contribute to the processing and integration of sensory information, formation of memories and oscillatory dysfunction in conditions such as epilepsy(*2*), schizophrenia(*3*) and Parkinson’s(*4*). Although many developments have been made in the ability of scientists and clinicians to capture neuronal activity data, no single approach is able to record all aspects of brain function. What is more, in the pre-clinical sphere the use of rodent disease models often requires long-term recordings, behavioural readouts and/or functional manipulations of neuronal firing. Each of these considerations limits the experimental methodology or recording technology usable for data acquisition. High-density, high-bandwidth systems based around silicon probes(*5*) or tetrode bundles(*6*) generally require tethered connections to amplifiers or digitisers, limiting the length and nature of recordings which can be performed and restricting the use of some behavioural experiments such as the continuous novel object recognition test(*7*), a highly sensitive experiment useful in models of schizophrenia. In contrast, fully wireless systems are limited by both the bandwidth available and the life of the implant battery(*8*), which will either require recharging, replacement or will limit the duration and frequency of recordings. Generally, a trade-off in wireless telemetry between bandwidth and battery must be made whereby the higher the bandwidth the shorter the batter life, or larger the battery pack must be(*9*). To compound this, even with a wireless telemeter the use of a stimulator (whether electrical or optical) usually necessitates further implants and tethers, again limiting the range of suitable environments, behavioural readouts, and length of recording sessions featuring the stimulus.

The study presented here takes a prototype wireless telemeter, with onboard optogenetic stimulator, which constantly charges via an inductive field. We develop an implantation strategy to completely contain the biopotential leads and optic fibre beneath the skin, meaning that rats have no protruding dental cement, tethers or wires. This approach opens up a wider range of possible behavioural readouts, allows the rats to present as ‘normal’ to other rodents, and reduces the incidence of side-effects arising from lesions or dehiscent wounds found in conventional implant strategies. Furthermore, the wireless charging of the telemeter allows for sample rates of up to 2KHz, allowing for the recording of higher-frequency events and greater resolution in sampling low-frequency activity.

## Methods

### Ethics

All procedures on rats were reviewed and agreed with the Bioethics Committees and AWERB at Aston University and were compliant with current UK Home Office guidelines and project and personal licences under the Animals (Scientific Procedures) Act 1986 and EU Directive 2010/63/EU.

### Surgery

All tools and equipment used in the virus injection and telemeter implantation surgery of the telemeter had been sterilised prior to starting the procedures via autoclave at 121ºC (Prestige Medical Classic Media Autoclave).

### Virus injections

PBS solution (Gibco 1X tablets) was made up in deionised water and then sterilised by autoclave. The virus stock solution (pAAV-Syn-ChR2(H134R)-GFP (AAV8), Addgene 58880-AAV8) was diluted in a 1:1 dilution factor with the sterile 0.1 PBS solution and stored on dry ice in preparation for the surgery. Rats of around 80g were used for the viral injection procedures. To begin the procedure, isoflurane anaesthesia was induced via a dedicated induction box before transferring the rat to the surgical rig and concentric mask (1.5-2.5% isoflurane delivered with 1.5L/m O_2_). The rat’s head was shaved, and the rat given an IP injection of Metacam and subcutaneous lidocaine over the skull. Once positioned in the stereotaxic frame and secured with ear bars, skin scrub was applied to the shaven area before making a midline incision in the skin on top of the skull with a sterile scalpel and clearing the incision site. Having located bregma, the coordinates of ML: -2.5, AP: -3 from bregma were marked and a hole in the skull was drilled with a conical burr. The prepared virus solution was defrosted and 1ul of the solution was slowly (over 10 minutes) injected into the cortex at DV: -1 using a Hamilton syringe. After 5 minutes dwell time and the slow removal (over 5 minutes) of the syringe, the head wound was sutured closed and a layer of cyanoacrylate skin glue was put on top, the nails were trimmed, and the rat was then put into a recovery cage on a heat pad to be monitored during its recovery.

### Telemeter implantation

When injected rats reached 175g their size was suitable for telemeter implantation. Inhalational anaesthesia was induced as above, analgesia provided as above and the rat positioned in a supine manner with a large concentric mask. The abdomen and head were shaved, with a further shaved path linking the two locations around the flank of the rat and between the shoulder blades to accommodate the sub-fascial route of the leads and fibre. Using a sterile swab, skin scrub was then applied to the shaven areas. With the rat supine, an incision was made along the midline of the abdomen in layers through the skin and then through the muscle. After sterilising the telemeter (AD Instruments TR58AB Optogenetics biopotential telemeter) during surgical prep by submerging it in Cidex for 20 minutes before thoroughly rinsing with sterile water, it was inserted into the visceral cavity and secured to the muscle wall using non dissolvable silk sutures through the suture tabs on the telemeter body. The muscle wall was then closed again using absorbable 5-0 vicryl sutures, leaving enough space for the optic fibre and biopotential leads to exit the visceral cavity through the incision. To create a subcutaneous path to the skull, blunt dissection was performed to lift the skin from the muscle. The biopotential leads and the optic fibre were then put through a steel trocar used to guide the fibre and biopotential leads up to the skull where an incision was made allowing the trocar to be pulled out through this incision, leaving a small amount of the fibre and biopotential leads protruding over the cranium. Sterile gauze was placed over the open wound on the abdomen before turning the rat over to a prone position and then secured in the stereotaxic frame. The skin on the skull was held open using clamps, bregma was then located, and the following coordinates were found, marked and then drilled:

ML: -2.5, AP: -3 (for the recording biopotential lead)

ML: 2.5, AP: -3 (for the reference biopotential lead)

ML: -2.5, AP: -4 (for the optic fibre)

EEG screws were inserted in the holes drilled for the biopotential leads and used to secure them in place by wrapping a section of the wire from the biopotential leads around their respective screws. The fibre was then inserted into the hole at a slanted angle (20º), advanced through the brain to sit below the EEG screws, and supported and secured by building up layers of dental cement around the fibre as well as the biopotential leads to create a low-profile dental cement cap on the skull. Extra screws (3-4) were also inserted into the skull in order to help secure the dental cement cap. When the dental cement cap was dry the wound was then sutured closed over the top of the cement cap using the vicryl sutures and cyanoacrylate skin glue. When the glue was dry, the rat was then returned to the supine position to suture the abdominal wound closed and skin glue was applied on top.

To reduce the chance of the rat scratching and exacerbating wounds, their nails were then trimmed before putting them in a recovery cage on a heat pad.

### Data acquisition

Once the rat had recovered from the surgery the rat’s home cage was placed on a Kaha Smartpad, connected to a PowerLab and Kaha Configurator (ADInstruments) connected to a Windows PC. Data were recorded using LabChart Lightning whilst the stimulation parameters of the telemeter were changed using Kaha ConfigSoft. Data were analysed post-hoc using LabChart8 and GraphPad Prism 10.

### Stimulation

ChR2-H134R(*10*) activity was induced via the onboard optogenetic stimulator, comprising a blue LED (470nm) feeding the implanted fibre-optic. Stimulus duration, frequency and intensity were controlled via the Kaha ConfigSoft system and Kaha Configurator. Frequencies from 1Hz to 50Hz were trialled and observed, with pulse widths and intensities altered to produce the clearest response for the minimum amount of stimulation.

### Telemeter explant and reconditioning

At the end of telemeter experiments rats were anaesthetised in an induction box with isoflurane before IM injection with xylazine (0.2ml/kg), pentobarbital (2ml/kg – 0.4ml/kg for older rats and 0.6-0.8ml/kg for younger rats) and ketamine (0.4ml/kg). To ensure that injectable anaesthesia was maintained rats were watched for several minutes outside of the anaesthetic chamber and frequent checks were performed on pedal reflex and breathing rate. After this, transcardial perfusion was then started by cutting through the skin and muscle layers over the ribcage and opening the abdomen just under the sternum, with care taken to avoid damage to the biopotential leads and the optic fibre. The diaphragm was then cut in order to gain access to the heart to place an 18-gauge butterfly needle through the left ventricle into the ascending aorta, then a cut was made into the right atrium whilst PBS solution (or, for histology, PBS followed by 4% formaldehyde in PBS) is pumped through for 5 minutes to flush out all remaining blood. An incision was then made over the skull in order to lift the cement cap off gently using a scalpel blade. The rat then was decapitated, and the brain removed and put into either artificial cerebrospinal fluid to then be used for further electrophysiology or formalin fixative solution for 24 hours before being transferred into PBS sucrose solution for histology. The telemeter was then removed from the rat by making careful incisions around the optic fibre and biopotential leads starting from the skull towards to abdomen where the sutures were removed from the muscle wall in order to lift the telemeter from the visceral cavity. The dental cement cap was dissolved using acetone.

Explanted telemeters were rinsed with deionised water and disinfected using Cidex for 20 minutes before being rinsed with water again to be ready for re-use. Fibres can be lengthened or replaced using zirconium connectors (ThorLabs CFLC230) and mating sleeves (ThorLabs ADAL1) with the identical fibre being used to replace the factory-suppled one (ThorLabs FP200URT).

## Results

### Subcutaneous implant strategy results in normally-presenting rats

A total of 20 male Wistar rats were implanted with the wireless optogenetic telemetry system in the manner described above (Figure 1). Refinements were made during the course of the experiments to improve fibre positioning and increase the robustness of the fibre especially around the neck musculature. The transition between the fascial tunnel through the flank and the head area is the most likely point of failure for the implanted fibre, and over the lifetime of the implant migration of the fibre deeper into the brain becomes more likely if a connector or ferrule is not used. To guard against fibre breakages, a biocompatible PTFE tubing (Bohlender 2284108) was added to later surgical implants which prevented any loss of animals due to fibre displacement or breakage.

**Figure 1:**
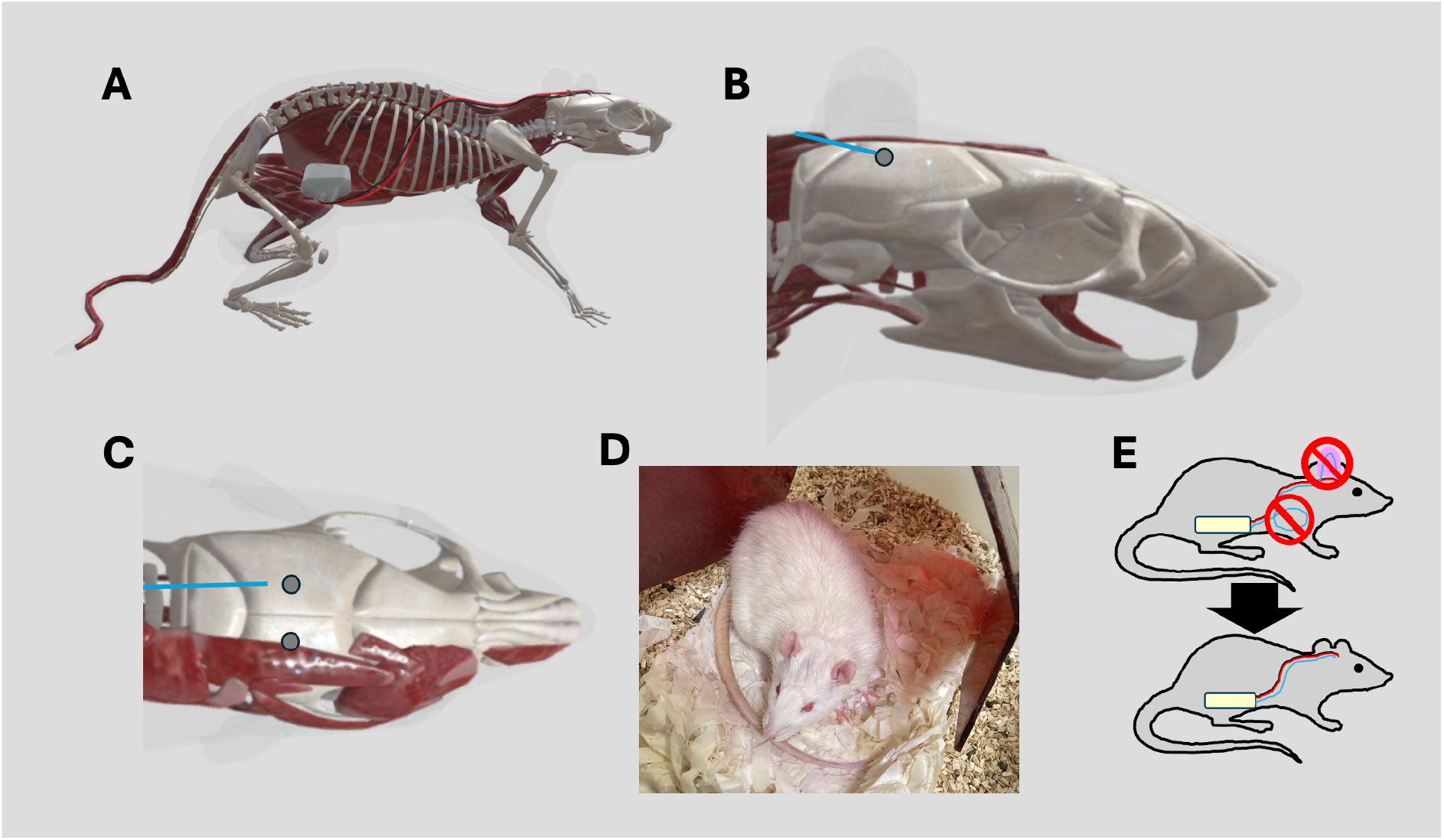
Implant strategy and location. A: Computer rendering of an adult (200g) rat with abdominal telemetry location and track of biopotential leads. Optogenetic fibre (not shown for clarity) can be coiled in the abdominal cavity, connectorized to split the fibre path or cut to size and run straight to the implant site. B-C: Side and plan view of implant site showing approximate location of EEG screws/depth electrodes and fibre placement. D: Rats once implanted and recovered present as normal. Male Wistar rat (250g) with onboard telemeter and sensorimotor implant of EEG screws and optic fibre shows only a small bulge on the top of the head, with skin and fur fully grown back. E: Overall implant strategy led to refinements in both headgear size and fibre location – cutting the fibre and refurbishing via connectors proved more sustainable and reliable than coiling the spare fibre within the abdomen.

Once implanted, rats presented as visibly normal without protrusions over the cranium or swelling at the abdomen (Figure 1E). Fluid and food intake were as with unimplanted animals and behaviour within the home cage and in open fields were normal throughout. During stimulation periods rats showed no visible signs of distress, twitching or aberrant behaviour.

As a further refinement and to aid the refurbishing and redeployment process of the telemeter the standard fibre was cut and replaced with an identical fibre-optic terminated with zirconium physical-contact ferrules, coupled via a zirconium sleeve. This did not result in any observable difference in implant performance or biocompatibility, with an average power-loss of 20% due to the connector (44mW/mm^2^ vs 55mW/mm^2^ at maximum power).

### Optogenetic stimulation of sensorimotor cortex entrains oscillations

In both acutely anaesthetised and freely-moving animals, a series of stimulation protocols were performed to assess the ability of the implanted fibre to produce reliable oscillatory activity. A broad expression strategy for the optogenetic construct and the H134R ChR2 variant was chosen to focus on ensemble activity rather than single-cell firing. An example trace is shown in Figure 2 – stimulation at 20Hz (20ms pulse duration, 80% total power output) clearly produces a reliable train of events in the raw EEG and in beta-band (13-30Hz) filtering. Activity was reliably entrained from 2Hz to 20Hz with both standard fibre configurations and with connectorized fibres (Figure 3). Data were filtered in real time into standard bands for canonical oscillations to ensure that the stimulus used was entraining the expected frequency of activity. No observable behavioural changes resulted from stimulus across the frequencies used.

**Figure 2:**
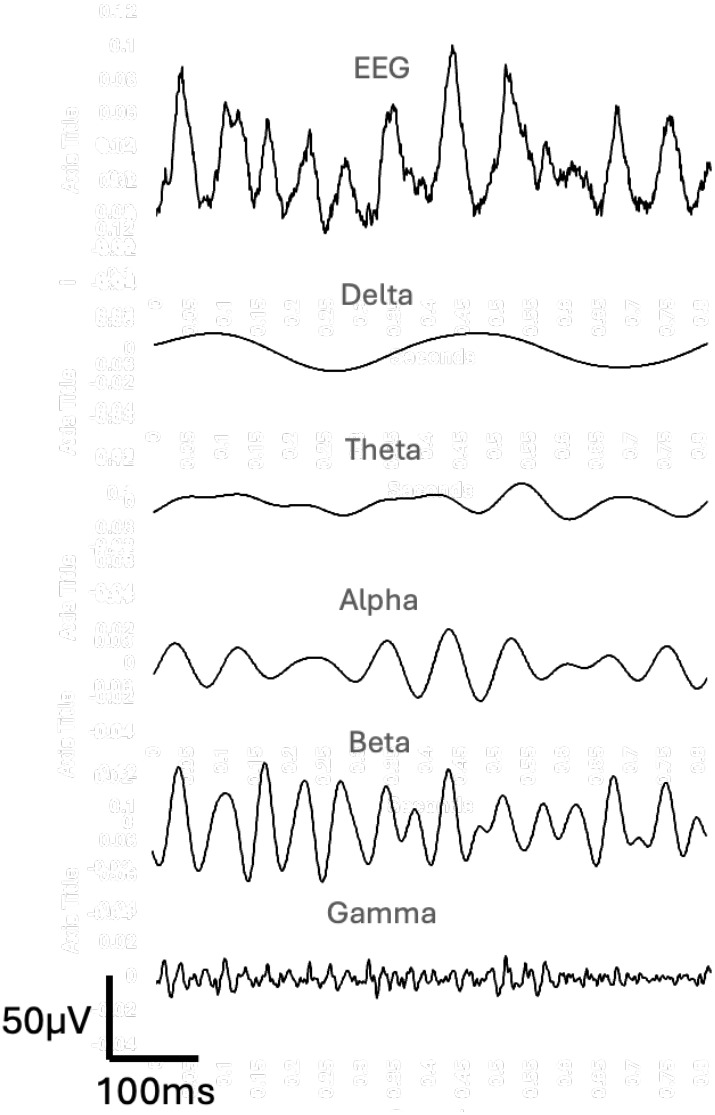
Example EEG shows optogenetic entrainment of oscillations at 20Hz. Raw EEG signal at top is filtered in real-time into multiple band-pass components reflecting standard ranges for Delta (1-4Hz) to Gamma (31-180Hz) band oscillations

**Figure 3:**
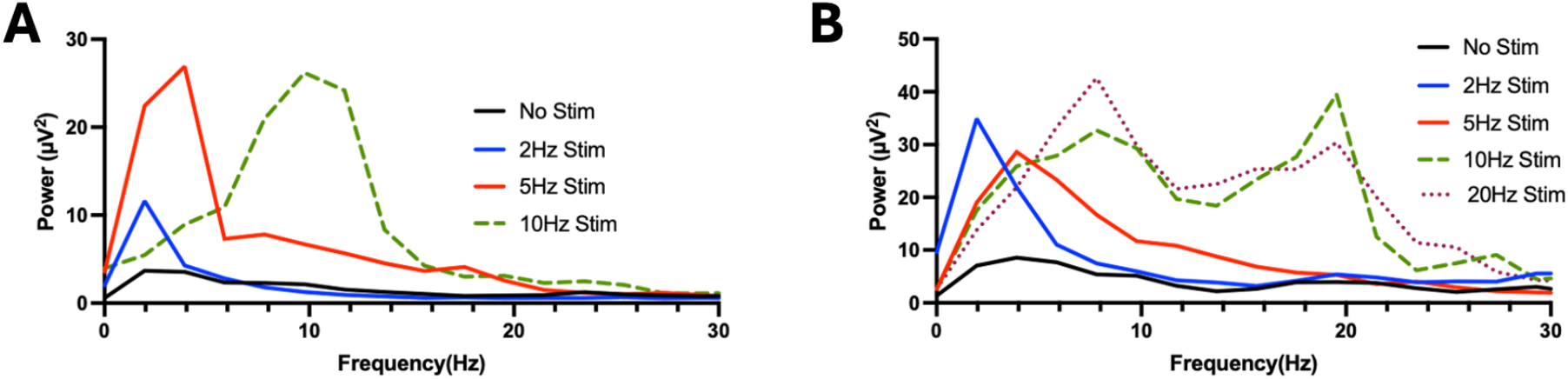
Stimulation from 2Hz to 20Hz allows entrainment of oscillations in both unmodified and connectorized implants. A: Factory-specification implant in a freely-moving adult Wistar rat shows clear increases in FFT power across stimulated frequencies. B: The same telemeter with a connectorized fibre (in a different rat) produces entrainment of a similar power. Most rats, whether implanted with connector-equipped telemeters or not, displayed harmonics (as seen in B at 10 and 20Hz) at some higher-frequency stimulation.

### Twenty Week implantation shows no loss of signal or entrainment ability

In a subset of rats (N=5) long-term implants of up to 6 months were performed. These were recorded throughout the implantation period without interruption, the implant protocol allowing for continuous data acquisition even during the cleaning/husbandry process which the animals experienced daily. Stimulation experiments were carried out at irregular intervals in clusters to assess the variability and reliability of stimulation throughout the lifetime of the implant. Figure 4 shows both the significant increase in FFT power driven by optogenetic stimulation (P <0.01, ratio paired t-test) and the stability of responses over 140 days of implantation. Experiments were performed in clusters to ensure that variance in the data were down to day-to-day differences rather than a progressive increase/decrease in response.

**Figure 4:**
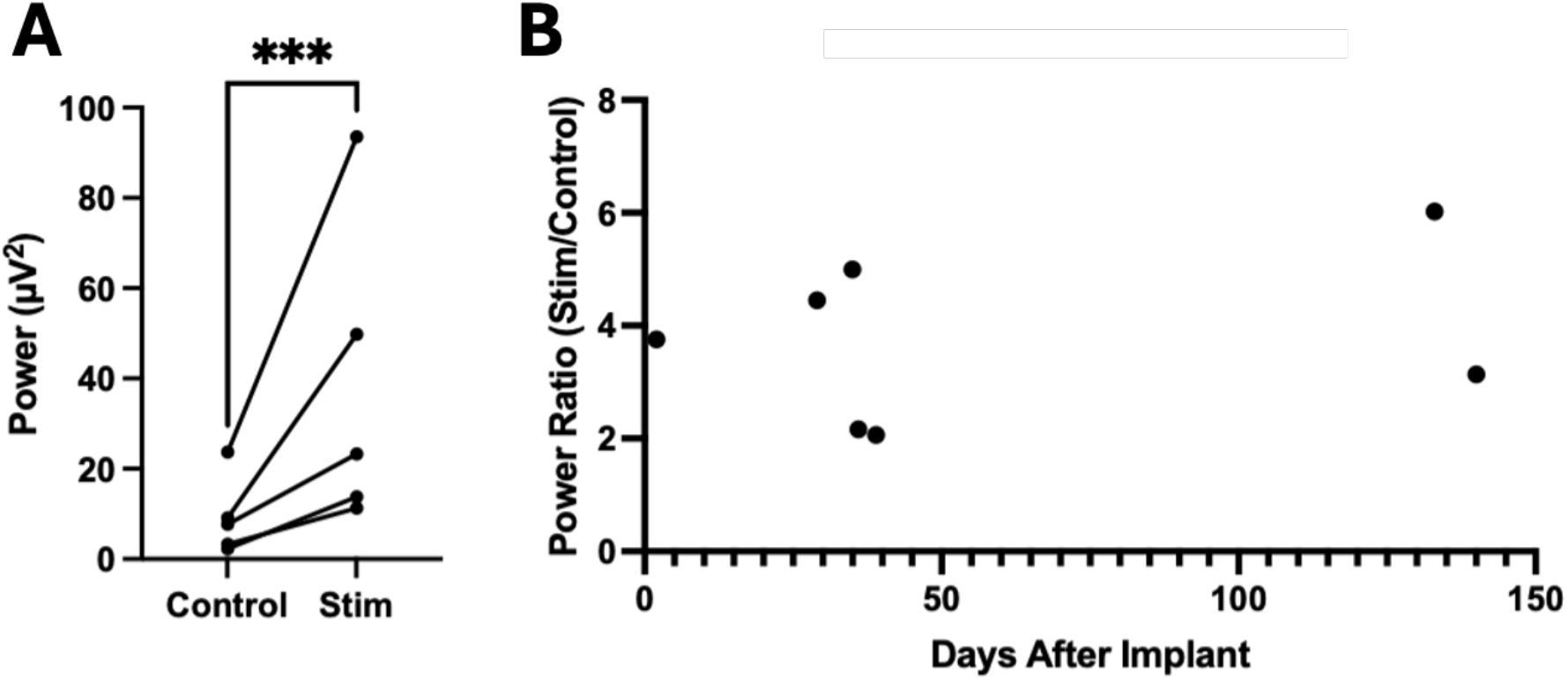
Optogenetic stimulation produces robust changes in oscillatory power and is stable long term. A: Paired data in freely-moving rats shows a significant increase in power under optogenetic stimulation. B: Experiments over 140 days show no significant changes in response to stimulation during the lifetime of the experiment.

## Discussion

Any attempt to record data from neurons, either from single cells or a whole brain, is subject to a range of compromises and considerations. Here we have presented a new approach to reducing some of those compromises via the combination of high-resolution telemetry with optogenetic stimulation, contained subcutaneously to allow for a wider range of behavioural experiments, fewer avenues for infection or implant breakdown and more naturalistic implanted animals. The implants are stable for six months (or beyond) with no loss of stimulation power or signal strength. Inserting the fibre at an angle, required for containment under the skin, will obviously limit the brain areas able to be targeted to largely cortical areas. However, the targeting of structures such as the basal ganglia is still possible if required via the creation of a small dental cement-based ‘cap’ during implantation providing adequate bend-radius to the implanted fibre. The primary advantages of the subcutaneous approach are to limit the opportunity for infection or wound dehiscence, to provide a more low-profile headstage for behavioural experiments, and to ensure the rat presents as ‘normal’ to other rodents for social behavioural tasks. To that end, this telemetry approach has a wide range of potential uses in fields such as epilepsy, schizophrenia(*11*) and autism studies(*12*). The increased sampling rate a/orded by the constantly wireless charging may be useful in investigating very high frequency oscillations seen in some of the above conditions(*13*).

It Is clear that the entrainment of oscillations in the cortex is shaped by the natural activity of the cells and circuits involved in the generation of physiological activity. It would be unreasonable to expect extrinsic stimulus to produce narrow, coherent activity in the specific frequencies stimulated in anything other than single units. Nevertheless, we have shown that a broad response with a peak approximating the primary stimulation frequency can be obtained, without obvious stimulation artefact or behavioural issues in freely-moving rats. We propose that this stimulated entrainment may be suitable for sleep studies and oscillopathies again including schizophrenia and epilepsy.

